# Whole transcriptome *in-silico* screening implicates cardiovascular and infectious disease in the mechanism of action underlying atypical antipsychotic side-effects

**DOI:** 10.1101/2020.04.09.033472

**Authors:** Yasaman Malekizadeh, Gareth Williams, Mark Kelson, David Whitfield, Jonathan Mill, David A Collier, Clive Ballard, Aaron R Jeffries, Byron Creese

## Abstract

**INTRODUCTION:** Stroke/thromboembolic events, infections and death are all significantly increased by antipsychotics in dementia but little is known about why they can be harmful. Using a novel application of a drug repurposing paradigm, we aimed to identify potential mechanisms underlying adverse events.

**METHOD:** Whole transcriptome signatures were generated for SH-SY5Y cells treated with amisulpride, risperidone and volinanserin using RNA-sequencing. Bioinformatic analysis was performed which scored the association between antipsychotic signatures and expression data from 415,252 samples in the NCBI GEO repository.

**RESULTS:** Atherosclerosis, venous thromboembolism and influenza NCBI GEO-derived samples scored positively against antipsychotic signatures. Pathways enriched in antipsychotic signatures were linked to the cardiovascular and immune systems (e.g. BDNF, PDGFR-beta, TNF, TGF-beta, selenoamino acid metabolism and influenza infection).

**CONCLUSION:** These findings for the first time mechanistically link antipsychotics to specific cardiovascular and infectious diseases which are known side effects of their use in dementia, providing new information to explain related adverse events.

## 1. Background

Atypical antipsychotics are a commonly used off-label treatment for agitation, aggression and psychosis in dementia. They are modestly effective but have a severe side effect profile which includes sedation, thromboembolic events, QTc prolongation, falls, fractures, infections, stroke and all-cause mortality [1,2]. The narrow margin of clinical benefit and the lack of alternative pharmacological agents makes investigation of drug safety a key priority. Antipsychotic therapeutic mechanism of action (MoA) is primarily via antagonism of serotonin receptor 2A (5-HT2A) and/or dopamine receptors 2 and 3 (D2/3) but many also have significant antihistaminergic, anticholinergic and antiadrenergic properties. It has long been hypothesized that this off target activity is a contributor to the side effect profile of antipsychotics in dementia [1,3–6]. It has also been suggested that generic mechanisms such as over sedation leading to dehydration, failure to clear the chest and inactivity may be key mediating mechanisms [1]. Therefore an important unanswered question is whether side effects are a primary result of perturbations to specific biological processes (e.g. cardiovascular biology, immune response) or secondary consequences of more general mechanisms like sedation. Understanding the answer to this question will help enormously in the future development of safer antipsychotics and inform the safer prescribing of existing agents.

High throughput *in-silico* screening approaches leveraging gene expression data may provide novel mechanistic insights into dementia-related side effects. Such approaches rest on the principle that transcriptional activity represents a useful surrogate for disease states and are widely used to triage compounds in drug repurposing studies (exemplified by the Connectivity Map, Cmap) [7–11]. A typical application would see a gene expression signature from a candidate disease screened against a compound expression database; negative scores indicating possible therapeutic benefits (i.e. evidence that the drug reverses the disease transcriptional signature). It follows that a positive score between a given compound and a condition which is a side effect of that compound would indicate a MoA which is linked to the condition. Thus a key advantage of this approach in the examination of drug side effects is a more direct biological link to human disease side effects without testing in humans.

In the present study, our aim was to determine whether transcriptional perturbations derived *in-vitro* could elucidate mechanisms underlying adverse effects of antipsychotic use in dementia. We generated gene expression signatures for three antipsychotics representing a range of mechanisms of action relevant to the current landscape of drug development and clinical use in dementia: amisulpride (primarily a D2/D3 antagonist), risperidone (primarily a 5HT2A/D2 antagonist) and volinanserin (highly selective 5HT2A inverse agonist) [12–14]. We then used a high-throughput bioinformatic scoring algorithm to test for association with human diseases. Specifically, we hypothesized that the antipsychotic signatures would be score positively with conditions and diseases related to known side effects of their use in dementia.

## 2. Materials and methods

### 2.1 Antipsychotics

The following antipsychotic concentrations were used, based on previously published doses [12,14–17]: 1μM amisulpride (catalogue number CAY14619, Cambridge Bioscience, UK), 100nM risperidone (catalogue number ab120393, Abcam, UK) and 10nM volinanserin (catalogue number CAY15936, Cambridge Bioscience, UK). Dimehtyl sulfoxide (DMSO) was used as the vehicle for all compounds.

### 2.2 Cell culture

SH-SY5Y human neuroblastoma cells (P13) were cultured in media (DMEM/F-12, GlutaMAX™ Supplement; catalogue number 11514436, Fisher Scientific, UK) containing filtered 10% fetal bovine serum (Gibco™ Fetal Bovine Serum, heat inactivated; catalogue number 11550356, Fisher Scientific, UK). Cells were maintained at 37°C, 5% CO_2_ and atmospheric O_2_ in a humidified incubator. Cells were seeded at a density of ~70% in 6-well plates the day before experimentation and grown in the same media. On the day of the experiment, cells were treated with filter sterilized media containing the antipsychotic compounds or vehicle at desired concentration for 24 hours. No cell death was observed at the drug doses tested. Four individual culture well replicates were collected for each compound and vehicle.

#### 2.3.1 RNA extraction

To preserve RNA in SH-SY5Y cells, media was removed and 500μl of Trizol (Invitrogen Trizol reagent; catalogue number 15596026, Fisher Scientific, UK) was applied to each well. Cells were mixed thoroughly with the reagent and collected for RNA extraction. RNA was purified using an RNA kit (Direct-zol™ RNA MiniPrep w/ Zymo-Spin™ IIC Columns (Capped); catalogue number R2052, Cambridge Bioscience, UK) as shown in the instruction manual and stored at −80°C. Following RNA purification, the concentration of RNA was measured by Qubit 3.0 Fluorometer using Qubit high sensitivity RNA kit (Qubit™ RNA HS Assay Kit; catalogue number Q32852, ThermoFisher Scientific, UK). The quality of purified RNA was tested using Agilent 2200 TapeStation system and RNA ScreenTape Assay (RNA ScreenTape; catalogue number 5067-5576, RNA ScreenTape Sample Buffer; catalogue number 5067-5577, Agilent, UK). The mean RIN value across all samples was 9.87 (minimum: 9.6, maximum: 10). RNA samples were diluted at the desired concentration for polyA-tail library preparation and sequencing.

#### 2.3.2 RNA Sequencing

Illumina HiSeq 2500 standard mode sequencing system was used to sequence RNA samples (poly-A tail library preparation, 125bp paired end, 20 million reads per sample). Quality control using FastQC was performed to remove low quality reads. To compare the expression profile of samples, STAR (version 2.6.1a) was employed to align the RNA reads to the reference human genome (hg38). To create and sort bam files, samtools (version 0.1.16) and to index and assign mapped reads to genomic features, featureCounts (version 1.6.1) were utilised.

### 2.4 Identification of differentially expression genes

To generate differentially expressed genes (DEGs), DESeq2 (version 1.16.1) was used which calculates and finds significant changes in samples based on negative binomial distribution. Statistical filtering based on the log2 of 1.5-fold change and a false discovery rate adjusted P-value (P_FDR_) <0.05 was used to generate the gene lists used in subsequent analysis. A 1.5-fold change cut off was applied so that genes perturbed due to off target (which may be relevant to side effects) as well as therapeutic actions of the compounds were captured.

### 2.5 High throughput screening of antipsychotic drug signatures against dementia-related side effects

To establish whether antipsychotic gene expression signatures were associated with gene expression of conditions representing known side-effects, we first conducted a high throughput *in-silico* screen against gene expression data from 415,252 human samples from 11,305 experimental series in the NCBI GEO repository using the Searchable Platform Independent Expression Database (SPIED, www.spied.org.uk) [18,19]. The SPIED tool facilitates querying of publicly available gene expression data from NCBI GEO with user-defined transcriptional signatures [8,9,11]. A major barrier to high throughput *in-silico* interrogation of human disease gene expression samples is that in many NCBI GEO series the case/control assignment of individual samples is not clear without manually curating the data (thus it is not practical to determine relative expression change across many hundreds or thousands of series). SPIED overcomes this by calculating an effective fold (EF) change at each probe in a sample, defined as the expression level of each individual array probe relative to the experimental series average [19].

In SPIED, association testing between the query antipsychotic signatures and NCBI GEO sample data is done via a Fisher Exact Test on 2×2 contingency table of up and down regulated genes. A score is assigned to each sample to reflect the relationship with antipsychotic expression. This score is defined as the sum of the number of genes perturbed in the same direction subtracted from the sum of number of genes perturbed in the opposite direction, divided by the total number of genes common to antipsychotic and sample profiles. Possible scores therefore range between −1 (all genes perturbed in the opposite direction) and 1 (all genes perturbed in the same direction), thus quantifying the relationship between an individual sample and query signature. If an NCBI disease series is associated with an antipsychotic then by definition individual samples within that series will positively score with the drug. This initial screen thus provides a first indication of association which can then be followed up. Specifically, highly scoring samples from NCBI GEO series assaying diseases or conditions of interest can then be manually assigned case/control status and tested for enrichment of positive scores among cases relative to controls. Thus, using SPIED, we followed the workflow described in detail in Williams (2013) [19] and broadly comprising of the following stages (graphically summarised in Figure 1):

1. Generate a statistically filtered list of differentially expressed genes for each antipsychotic (described in Section 2.4).
2. Use SPIED to screen each antipsychotic signature against all human gene expression micro-array data in the NCBI GEO repository. The resulting SPEID output is a ‘longlist’ of the 500 top scoring NCBI GEO samples with a statistically significant (adjusted P-value 0.05/11,305 NCBI GEO series= P<4.42*10^-6^) score (either positive or negative). The list was then manually curated to shortlist samples from NCBI GEO series meeting the following criteria:

a. Sample is from a series assaying one of the following disease areas relevant to side effects of antipsychotic use in dementia: thromboembolic events, stroke, bone density/osteoporosis (relevant to fractures), pneumonia and other respiratory infections, urinary tract infections and atherosclerosis/coronary artery disease.
b. Case/control design.
3. Manually annotate every sample in each shortlisted series as case or control according to their designation in NCBI GEO.
4. Test for enrichment of positive scores among cases relative to controls in each series using Fisher test. Given the correlation between the three antipsychotic signatures, a Bonferroni correction of 0.05/N shortlisted series was applied.

**Figure 1.**
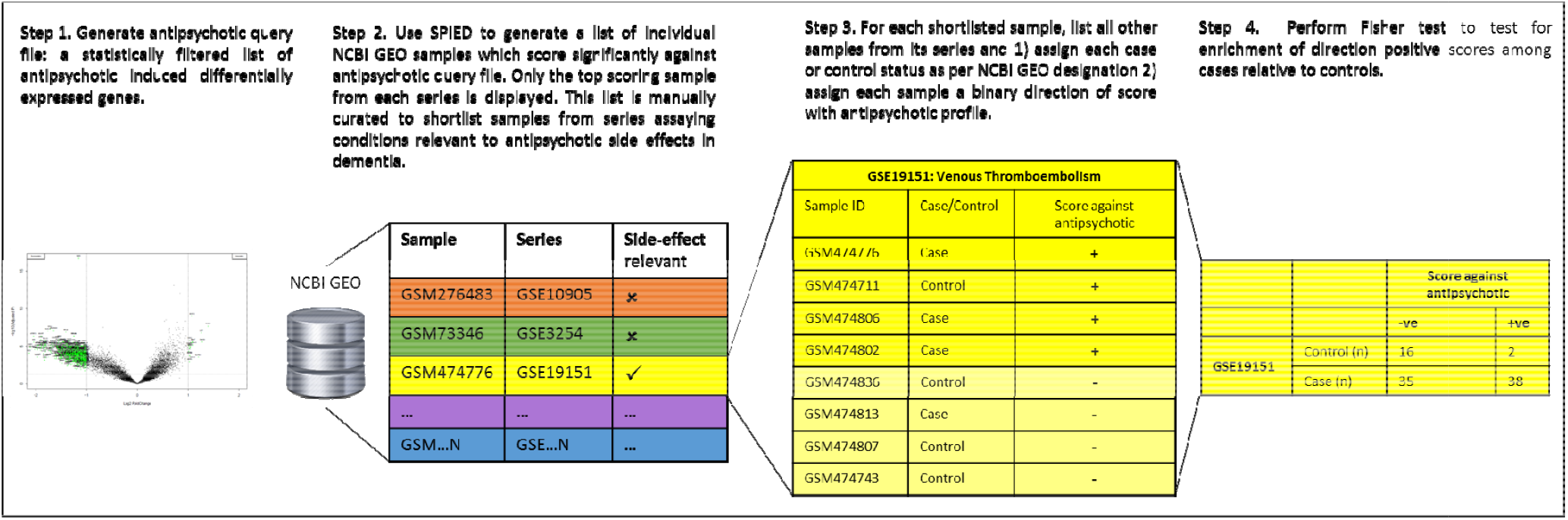
Graphical representation of SPIED screening method

## 3. Results

### 3.1 Differentially expressed genes

In total, 10,841 genes were detected and used for differential gene expression analysis. Gene expression level and bidirectional distribution pattern of expression associated with each antipsychotic is illustrated in the volcano plots presented in Figure 2. Treatment of cells with volinanserin, amisulpride and risperidone resulted in the activation of 2267 (1749 down-regulated and 518 up-regulated), 1026 (922 down-regulated and 104 up-regulated) and 809 (756 down-regulated and 53 up-regulated) genes, respectively (Fig 2, Supplementary Tables S1-S3). The three antipsychotic signatures were positively correlated with each other (amisulpride vs risperidone, Spearman test: r_s_= 0.76, amisulpride vs. volinanserin: r_s_ =0.88, risperidone vs volinanserin: r_s_=0.66).

**Figure 2.**
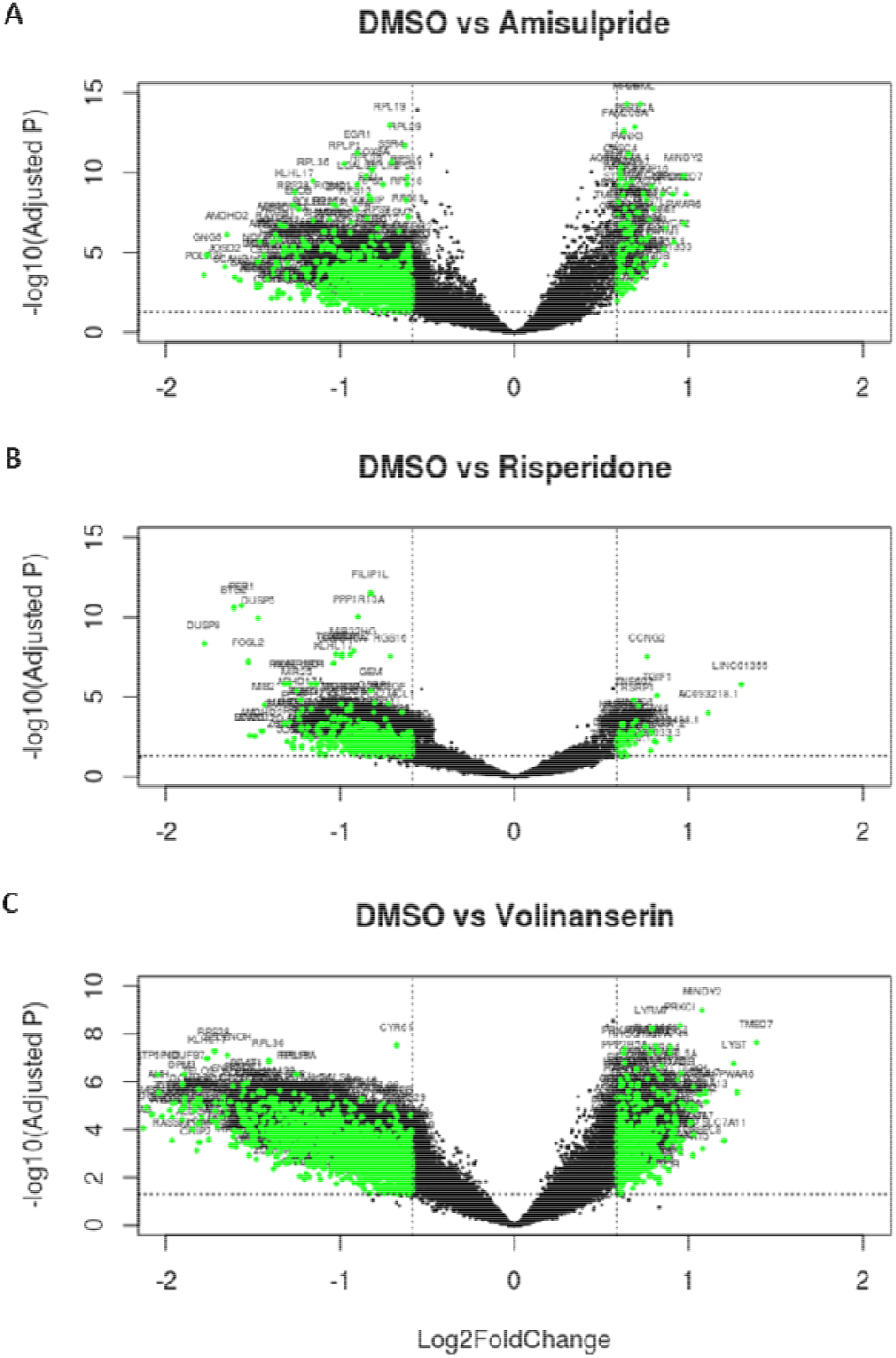
Volcano plots illustrating differentially expressed genes for amisulpride, risperidone and volinanserin vs. DMSO. Dotted horizontal lines mark adjusted p-value threshold of 0.05; dotted vertical lines mark log 1.5 fold change threshold. Green markers indicate statistically significantly differentially expressed genes with > +/-1.5 fold change.

### 3.2 Association between antipsychotic and dementia-related side effects

Each antipsychotic signature was screened against the NCBI GEO repository using SPIED (Step 2, Figure 1). As this is a high-throughput screen, we focused on the top 500 statistically significant (Bonferroni adjusted P-value 0.05/11,305 NCBI GEO series: P<4.42*10^-6^) scoring samples identified by SPIED for each drug. Of the 1500 total antipsychotic-sample scores identified by SPEID, 817 were statistically significantly associated with at least two antipsychotics, leaving 683 unique samples in the long list. This list of samples along with associated scores, p-value and number of overlapping genes is shown in Supplementary Table S4. Of these 683 unique samples, 18 were from series which assayed diseases/conditions relevant to side effects of antipsychotics in dementia (Step 3, Figure 1). Twelve of these were excluded as they were not casecontrol designs (meaning testing for association between the score in individual samples and case/control status is not possible). Thus six series were taken forward for further analysis: GSE13850 and GSE2208 (bone density), GSE23746 (atherosclerosis), GSE19151 (venous thromboembolism, VTE), GSE7638 (coronary artery disease, CAD), GSE17156 (respiratory infection, containing three conditions: influenza, rhinovirus and respiratory syncytial virus which were analysed separately in this analysis). Individual sample level data showing the distributions of cases and controls in each series and their associated scores and p-values are shown in Supplementary Tables S5 to S27 (Step 4, Figure 1).

Table 1 shows that atherosclerosis cases (GSE23746) were enriched for positive scores for all three antipsychotics (Fisher exact test amisulpride, p=0.002; risperidone, p=6.98×10^-5^, volinanserin, p=5.5×10^-3^). VTE cases (GSE19151) were enriched for positive scores for risperidone (p=8.13×10^-7^) and volinanserin (p=0.002). Finally influenza cases (GSE7638) were enriched for positive scores for amisulpride (p=0.002).

**Table 1.**
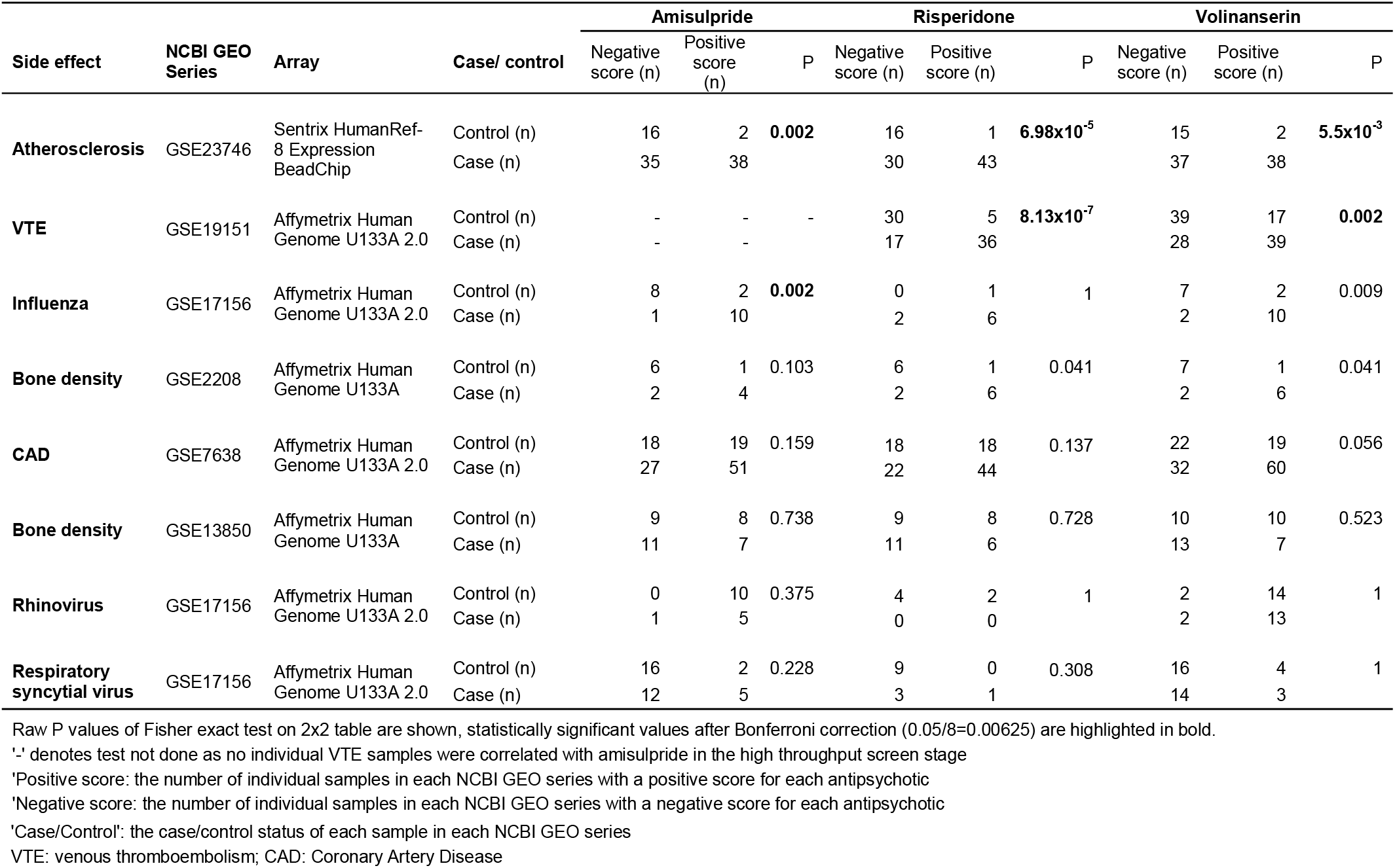
Association between antipsychotic and side effect gene expression profiles

| Side effect | NCBI GEO Series | Array | Case/ control | Amisulpride |  |  | Risperidone |  |  | Volinanserin |  |  |
| --- | --- | --- | --- | --- | --- | --- | --- | --- | --- | --- | --- | --- |
|  |  |  |  | Negative score (n) | Positive score (n) | P | Negative score (n) | Positive score (n) | P | Negative score (n) | Positive score (n) | P |
| Atherosclerosis | GSE23746 | Sentrix HumanRef-8 Expression BeadChip | Control (n) | 16 | 2 | **0.002** | 16 | 1 | **6.98x10<sup>-5</sup>** | 15 | 2 | **5.5x10<sup>-3</sup>** |
|  |  |  | Case (n) | 35 | 38 |  | 30 | 43 |  | 37 | 38 |  |
| VTE | GSE19151 | Affymetrix Human Genome U133A 2.0 | Control (n) | - | - | - | 30 | 5 | **8.13x10<sup>-7</sup>** | 39 | 17 | **0.002** |
|  |  |  | Case (n) | - | - |  | 17 | 36 |  | 28 | 39 |  |
| Influenza | GSE17156 | Affymetrix Human Genome U133A 2.0 | Control (n) | 8 | 2 | **0.002** | 0 | 1 | 1 | 7 | 2 | 0.009 |
|  |  |  | Case (n) | 1 | 10 |  | 2 | 6 |  | 2 | 10 |  |
| Bone density | GSE2208 | Affymetrix Human Genome U133A | Control (n) | 6 | 1 | 0.103 | 6 | 1 | 0.041 | 7 | 1 | 0.041 |
|  |  |  | Case (n) | 2 | 4 |  | 2 | 6 |  | 2 | 6 |  |
| CAD | GSE7638 | Affymetrix Human Genome U133A 2.0 | Control (n) | 18 | 19 | 0.159 | 18 | 18 | 0.137 | 22 | 19 | 0.056 |
|  |  |  | Case (n) | 27 | 51 |  | 22 | 44 |  | 32 | 60 |  |
| Bone density | GSE13850 | Affymetrix Human Genome U133A | Control (n) | 9 | 8 | 0.738 | 9 | 8 | 0.728 | 10 | 10 | 0.523 |
|  |  |  | Case (n) | 11 | 7 |  | 11 | 6 |  | 13 | 7 |  |
| Rhinovirus | GSE17156 | Affymetrix Human Genome U133A 2.0 | Control (n) | 0 | 10 | 0.375 | 4 | 2 | 1 | 2 | 14 | 1 |
|  |  |  | Case (n) | 1 | 5 |  | 0 | 0 |  | 2 | 13 |  |
| Respiratory syncytial virus | GSE17156 | Affymetrix Human Genome U133A 2.0 | Control (n) | 16 | 2 | 0.228 | 9 | 0 | 0.308 | 16 | 4 | 1 |
|  |  |  | Case (n) | 12 | 5 |  | 3 | 1 |  | 14 | 3 |  |

Raw P values of Fisher exact test on 2x2 table are shown, statistically significant values after Bonferroni correction ( $0.05/8=0.00625$ ) are highlighted in bold.
'-' denotes test not done as no individual VTE samples were correlated with amisulpride in the high throughput screen stage
'Positive score': the number of individual samples in each NCBI GEO series with a positive score for each antipsychotic
'Negative score': the number of individual samples in each NCBI GEO series with a negative score for each antipsychotic
'Case/Control': the case/control status of each sample in each NCBI GEO series
VTE: venous thromboembolism; CAD: Coronary Artery Disease

#### 3.2.1 Pathway analysis

Pathway analysis was then performed to elucidate more specific biological mechanisms underlying the reported associations. As this study is focused on side effects rather than therapeutic action, a pruned gene list for each antipsychotic was created; this comprised only of genes which were also differentially expressed in cases relative to controls in the series in Table 1. Thus the first step was to create a list of DEGs for atherosclerosis, VTE and influenza. This was done using the NCBI GEO analyser tool using a P_FDR_<0.05 threshold (gene lists for each signature are shown in Supplementary Tables S28-S30). DEGs in each antipsychotic signature which were not also present in any of the side effects signatures were excluded, creating three pruned gene lists.

For amisulpride, risperidone and volinanserin, query lists for pathway analysis comprised of 547, 435 and 1218 genes respectively (i.e. those genes overlapping with atherosclerosis, VTE or influenza). Genes in each of these three pruned antipsychotic lists were ranked in descending order by the log-fold change associated with the antipsychotic and tested for enrichment using the g:Profiler tool, which is well suited to pruned lists [20]. Gene set enrichment analyses included the following Gene Ontology (GO) and biological pathway sources: GO molecular function (MF), GO cellular components (CC), GO biological processes (BP), KEGG, REACTOME and WikiPathways. Any annotations not curated manually (therefore being less reliable) were excluded. g:Profiler’s multiple testing correction was applied (known as ‘g:SCS’ and developed specifically for pathway analysis). A g:SCS-adjusted P-value threshold of 0.05 was used [21]. Outputs were filtered to exclude pathway gene sets with <10 or >200 genes and with <3 overlapping genes in the input list.

##### 3.2.1.1 Biological pathways

Genes from 39, 23 and 44 GO terms and pathways were enriched in amisulpride, risperidone and volinanserin respectively (Figure 3, with detailed results in Supplementary Table S31).

**Figure 3.**
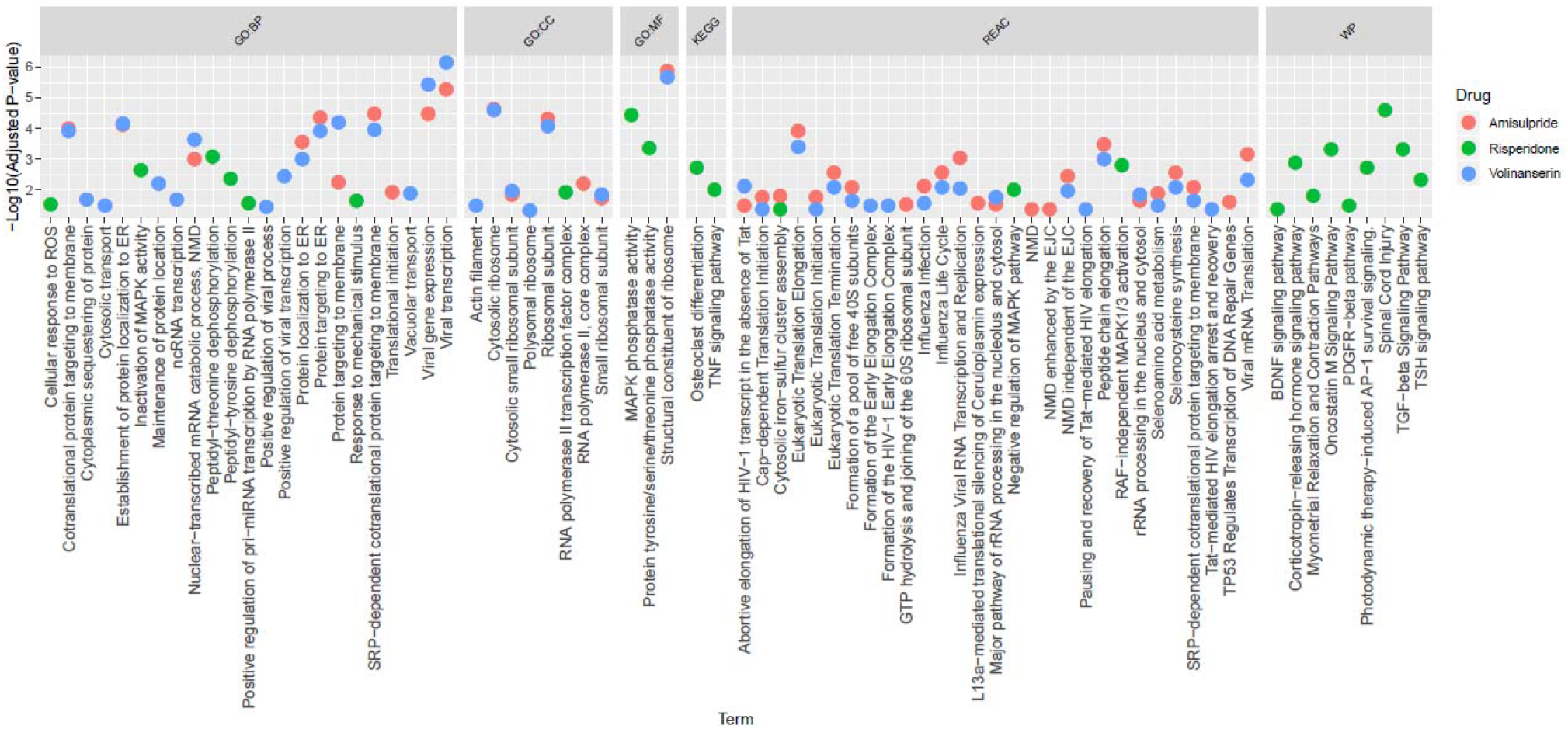
Plot of GO terms and pathways statistically significantly enriched in amisulpride, risperidone and volinanserin Abbreviations: GO, Gene Ontology; BP: Biological Processes; CC: Cellular Component; MF: Molecular Function; KEGG, Kyoto Encyclopedia of Genes and Genomes; REAC, REACTOME; WP, WikiPathways; NMD, Nonsense-mediated Decay; ER, Endoplasmic Reticulum; MAPK, Mitogen-activated Protein Kinase; EJC, Exon Junction Complex; GTP, Guanosine-5’-triphosphate; ROS, Reactive Oxygen Species; TNF, Tumor Necrosis Factor; RAF, Rapidly Accelerated Fibrosarcoma; TGF, Transforming Growth Factor; TSH, Thyroid Stimulating Hormone; PDGFR, Platelet Derived Growth Factor Receptor; BDNF, Brain-Derived Neurotrophic Factor.

Twenty-three and 21 Reactome pathways were enriched in amisulpride and volinanserin respectively. A number of these related to infectious disease pathways (e.g. viral mRNA transcription: volinanserin, g:SCS adjusted P=6.75x^-4^, amisulpride P=0.005; influenza life cycle: amisulpride, P=0.003, volinanserin, P=0.009). Two pathways linked to the essential amino acid selenium were also enriched in both amisulpride and volinanserin: selenocysteine synthesis (amisulpride, P=0.003, volinanserin, P=0.009) and selenoamino acid metabolism (amisulpride, P=0.01, volinanserin, P=0.03).

For risperidone, 14 pathways across the KEGG (n=2), Reactome (n=3) and Wikipathways (n=9) databases were identified. The Reactome pathways were linked to MAPK (RAF-independent MAPK1/3 activation, P=0.002; Negative regulation of MAPK pathway, P=0.01). KEGG and WikiPathways enriched in risperidone were linked to cell growth/differentiation, with some growth factor pathways linked to the cardiovascular system and inflammation: brain derived neurotrophic factor (BDNF) signalling pathway, P=0.045; platelet derived growth factor receptor (PDGFR)-beta signalling, P=0.034; osteoclast differentiation, P=0.002; inflammation; oncostatin M signalling, P=0.0005; transcription necrosis factor (TNF) signalling pathway, P=0.01; transforming growth factor (TGF) beta signalling, P=4.6x^-4^.

##### 3.2.1.2 GO terms

All GO terms enriched in the three antipsychotic lists are shown in Figure 3, with detailed results in Supplementary Table S31. Removing redundant terms using Revigo [22] showed that the amisulpride gene list was primarily enriched for GO terms related to viral transcription (P=1.29x^-6^), SRP-dependent co-translational protein targeting to membrane (P=3.43E-05), cystolic ribosome (P=2.2x^-5^) and structural constituent of ribosome (P=1.29x^-6^).

Risperidone was enriched for terms relating to peptidyl-threonine dephosphorylation (P=8.43x^-4^), response to mechanical stimulus (P=0.02), positive regulation of pri-miRNA transcription from RNA polymerase II promotor (P=0.03), RNA polymerase II transcription factor complex (P=0.01), MAP kinase phosphatase activity (P=3.5x^-5^).

Volinanserin was enriched for viral transcription (P=6.94x^-7^), SRP-dependent co-translational protein targeting to membrane (P=1.07E-04), cystolic ribosome (P=2.46x^-5^) and structural constituent of ribosome (P=2.03x^-6^).

## 4. Discussion

This study aimed to elucidate mechanisms underlying side effects associated with antipsychotic use in dementia. To our knowledge we provide the first evidence mechanistically linking antipsychotics with specific cardiovascular and infectious diseases which are common side effects of their use in dementia. Supporting our hypothesis, the initial high throughput screen identified three conditions related to known side-effects which were associated with the antipsychotics; atherosclerosis cases were enriched for positive scores with all three antipsychotics, venous thromboembolism cases were enriched with positive scores for risperidone and volinanserin, and influenza cases were enriched with positive scores for amisulpride. Supplementing these drug-disease associations, a number of biological pathways related to cardiovascular biology, infectious disease and inflammation/immune system were enriched across antipsychotic signatures. These findings suggest specific cardiovascular and immune processes may underlie some harmful effects of antipsychotics and for the first time provide a number of candidates which can now be prioritised for further investigation.

Notable pathways enriched in risperidone include BDNF, PDGFR-beta, TNF and TGF-beta signalling. Findings from previous *in-vitro* and *in-vivo* studies strongly implicate PDGFR-beta in atherosclerosis and cardiovascular disease, providing a possible mechanism to explain the positive association between the three antipsychotics and atherosclerosis and VTE observed in this study [23]. Similarly, BDNF also plays a role in the cardiovascular disease (as well as neuroplasticity and development) [24,25] and is expressed in a variety of blood cells, the heart and vasculature [26]. It is also noteworthy that previous studies have demonstrated that part of risperidone’s pro-cognitive therapeutic mechanism of action may be via BDNF [27]. It is evident from our findings that more work must be done to untangle this complex element of antipsychotic MoA, where BDNF is plausibly related to both beneficial and detrimental effects of antipsychotics, which is highly relevant to dementia where the margin between clinical benefit and harm is so narrow. Two pathways linked to the essential amino acid selenium were enriched in amisulpride and volinanserin. Selenium plays a role in preventing oxidative stress and has been widely linked in observational studies to cardiovascular disease and atherosclerosis [28]. Moreover, one study in patients with schizophrenia implicated selenium deficiency in the adverse cardiac effects of clozapine, though it was not clear whether the deficiency was caused by the drug or the schizophrenia itself [29]. Our findings bring greater clarity to this previous work by providing evidence that antipsychotics directly act on selenium pathways. This has particular relevance to neurodegeneration where selenium deficiency in Alzheimer’s disease brain tissue has been observed and is hypothesised to play a role in cardiovascular side effects in Parkinson’s disease [30,31]. Our findings provide a clear indication for prioritising study of selenium deficiency and its interaction with antipsychotics in people with neurodegenerative disease in order to understand if it may be a clinically useful marker.

Infectious disease and immune pathways were also enriched across all three antipsychotics. These included a range of viral and influenza-linked GO terms in amisulpride and volinanserin, and TNF and TGF-beta in risperidone. Consistent with this, a recent study showed a considerable global suppression of immune response in mice treated with risperidone, indicated by reduction in a number of cytokines during treatment [32]. Our findings suggest that this impact extends to other antipsychotics and so underscore the need to prioritise investigation of immune response in people with dementia. They also suggest that susceptibility to infection associated with antipsychotics is not solely secondary to more general effects of antipsychotics like sedation-induced inactivity or failure to clear the chest.

Although more work needs to be done to build on the candidate mechanisms highlighted in this study, their initial identification is an important step which could ultimately have important implications for clinical decision making. For example, the incorporation of more formal cardiovascular history screening, with a particular focus on thrombosis risk or selenium deficiency, into clinical decision making could result in greater harm reduction.

We note that there were differences in the pathways enriched between antipsychotics however it would not be appropriate to draw direct comparison between them at the specific pathway level or interpret differences as clinically relevant. This is because these experiments were conducted *in-vitro,* so cellular responses will be affected by dosing and duration of exposure to each compound, similarly, equivalent doses and bioavailability of drugs in humans will differ. At a broader level however, it is worth noting that associations between antipsychotics and side-effects, and enrichment of relevant biological pathways were observed across all compounds, despite their differing MoAs. Further comparison in different biological models, including those where ageing and frailty can be incorporated, and epidemiological studies is now warranted [33]. This line of investigation could have important implications for Alzheimer’s disease, Parkinson’s disease, and elderly people with schizophrenia where clinical trials of amisulpride and pimavanserin (a highly selective 5-HT2A inverse agonist) have recently been published and more antipsychotic-like drugs are in development [2,34–36].

The overall trend towards downregulation of genes in this experiment is also worth comment. This pattern was particularly notable in risperidone, where 53 genes were upregulated and 756 were downregulated. However, although notable this is not without precedent. One study, with a similar design, which treated SK-N-SH neuroblastoma cell lines with risperidone for 24 hours showed 80% of genes were downregulated in analysis of microarray data [12].

With regard to limitations, the design and analysis of this study follows the same principles as Cmap and therefore the same caveats apply. These include the comparison between cell line-derived signatures and human studies, specifically that it would be premature to draw concrete conclusions on the clinical profile of compounds based on these data alone. However, as with Cmap, the trade-off is an experimental design which provides a high throughput low cost screen, analogous to a drug repurposing experiment where thousands of licensed compounds are triaged against a single disease signature. Similarly, in this study, screening three antipsychotic signatures against thousands of diseases showed that mechanisms underlying venous thromboembolism, atherosclerosis and infection may be relevant to the side effect profiles of antipsychotics, providing a clear rationale for prioritising their investigation in different biological models and epidemiological studies. In doing so, this study also represents an important step towards safety screening for compounds in development of neuropsychiatric symptoms in Alzheimer’s disease.

In summary, this study highlights molecular level links between cardiovascular and infectious diseases and antipsychotics, which in future may have important implications for use of existing compounds in clinical practice and the development of safer drugs for dementia in the future.

## ACKNOWLEDGEMENTS

This work was generously supported by the Wellcome Trust Institutional Strategic Support Award (204909/Z/16/Z) and in part through the MRC Proximity to Discovery: Industry Engagement Fund (External Collaboration, Innovation and Entrepreneurism: Translational Medicine in Exeter 2 (EXCITEME2) ref. MC_PC_16072.

